# Single cell transcriptomics-level Cytokine Activity Prediction and Estimation (SCAPE)

**DOI:** 10.1101/2023.10.17.562739

**Authors:** Azka Javaid, H. Robert Frost

## Abstract

Cytokine interaction activity modeling is a pressing problem since uncontrolled cytokine influx is at fault in a variety of medical conditions, including viral infections like COVID19, and cancer. Accurate knowledge of cytokine activity levels can be leveraged to provide tailored treatment recommendations based on individual patients’ transcriptomics data. Here, we describe a novel method named Single cell transcriptomics-level Cytokine Activity Prediction and Estimation (SCAPE) that can predict cell-level cytokine activity from scRNA-seq data. SCAPE generates activity estimates using cytokine-specific gene sets constructed using information from the CytoSig and Reactome databases and scored with a modified version of the Variance-adjusted Mahalanobis (VAM) method adjusted for negative weights. We validate SCAPE using both simulated and real single cell RNA-sequencing (scRNA-seq) data. For the simulation study, we perturb real scRNA-seq data to reflect the expected stimulation signature of up to 41 cytokines, including chemokines, interleukins and growth factors. For the real data evaluation, we use publicly accessible scRNA-seq data that captures cytokine stimulation and blockade experiment conditions and a COVID19 transcriptomics data. As demonstrated by these evaluations, our approach can accurately estimate cell-level cytokine activity from scRNA-seq data. Our model has the potential to be incorporated in clinical settings as a way to estimate cytokine signaling for different cell populations within an impacted tissue sample.

## 1. Introduction

Single cell RNA-sequencing (scRNA-seq) technologies have transformed the study of transcriptomics from bulk tissue assays to single cell-level resolution. The cell expression profiles provided by scRNA-seq data enable examination of complex tissues that contain a mixture of cell types, phenotypes, and cell-cell interactions, e.g., the tumor microenvironment (TME). Within the scRNA-seq-based expression modeling space, cytokine activity prediction is an important topic since cytokines are one of the central agents in cell signaling and have been implicated in a number of medical conditions, including COVID-19, and cancer, due to acute, inflammatory distress and pervasive tissue damage. Cytokines, like interleukin-6 (IL6), for example, are one of the key drivers of the chimeric antigen receptor (CAR) T cell therapy-induced cytokine storm, which is a toxic side effect of therapy and results in physiological manifestations of fever, hypotension and respiratory insufficiency.^1^ Existing interaction detection methods, such as SingleCellSignalR,^2^ iTALK,^3^ CellPhoneDB^4^ and DiSiR (Dimer Signal Receptor Analysis),^5^ have two key limitations that impact their utility for cytokine activity prediction: 1) they operate at the level of cell types or clusters rather than at the level of individual cells, and 2) they aim to quantify interaction potential vs. interaction status. A cluster-level ligand-receptor interaction analysis ignores signaling heterogeneity within the analyzed groups. A population level approach can also mask important intercellular variations that are clinically relevant and can be used to resolve mixtures of cells into individual stromal, immune and cancer cells. Accurately characterizing this intercellular variation is important for the development of combination therapies as opposed to monotherapies, which can precisely target individual cell subpopulations.^6^

Existing cell-cell interaction methods do not differentiate between interaction potential (i.e., the receptor is present on the cell surface as inferred by elevated expression) and interaction activity (i.e., the receptor is present and bound by the cognate ligand). This limitation relates to a broader weakness of existing interaction estimation strategies in that they make the assumption that gene expression reflects protein abundance and that protein abundance alone is sufficient to predict protein-protein interaction strength. These assumptions do not account for the expression of binding factors, including transcription factors, related to each protein and, moreover, expression of signaling partners for each protein within real, diverse biological contexts (e.g., inflammatory environments). While methods that incorporate expression from genes downstream of the receptor, such as NicheNet,^7^ do exist, they depend on a definition of receiver and sender subpopulations to fully characterize cell signaling. Prior specification of receiver and sender cell populations to locate the ligand and/or the receptor origin is often challenged since in real-world signaling scenarios, the location of the originating ligand is ambiguous given that the ligand may simultaneously originate from several different groups of cells. Specifically, NicheNet is limited for the task of cytokine activity estimation since it results in a ranked list of ligand activity according to the ligand’s ability to predict observed differentially expressed genes compared to the background expressed genes as opposed to individual single-cell level ligand activity predictions.

There are several challenges associated with estimating cytokine activity at the single cell level. First, scRNA-seq data is extremely sparse (i.e., a large fraction of observed counts are zero) and has high levels of technical noise. As noted, this sparsity can unexpectedly influence expression correlation-based continuous interaction scoring methods (i.e., methods that score ligand-receptor pairs using data that contains multiple measurements for two population groups), ultimately leading to correlation values that quantify sparsity^8^ as opposed to biological associations between a ligand-receptor pair. A second challenge related to cytokine activity estimation is that cytokines are often small secreted proteins with redundant and pleiotropic functionality that cannot be accurately quantified by conventional abundance estimation technologies such as CITE-seq (Cellular Indexing of Transcriptomes and Epitopes by Sequencing).^9^

Here we propose the Single cell transcriptomics-level Cytokine Activity Prediction and Estimation (SCAPE) method that leverages a weighted gene set scoring mechanism to predict the activity of 41 human cytokines using scRNA-seq data. We validate the SCAPE method using both a simulation framework and real stimulated and blockade experiment-based scRNA-seq data as well as the Liao et al. COVID19 dataset.^10^ As compared with existing ligand activity prediction techniques, our method generates continuous, cell-level cytokine activity estimates using the expression of genes with experimental evidence of upregulation following cytokine stimulation. In addition, since our method depends on expression from genes downstream of the ligand as opposed to the individual ligand expression, it does not entail specification of the receiver and sender subpopulations. Our approach for predicting cell-level cytokine activity has the potential to be leveraged in hospital/clinical settings that have access to single-cell transcriptomics facilities for analyzing patient samples. Physicians can additionally compute relative changes in cytokine activity levels via multiple scRNA-seq samples to develop a more nuanced understanding of patient disease.^6^ This data on predicted activity can then be leveraged to identify compensatory cytokines, which have a better safety profile when treating immunotherapeutic side effects of conditions such as cancer.

## 2. Methods

### 2.1. SCAPE method for scRNA-seq data

The Single cell transcriptomics-level Cytokine Activity Prediction and Estimation (SCAPE) method estimates the cell-level activity of target cytokines using scRNA-seq data. Figure 1 provides a high-level overview of the approach. An R package implementing both the CytoSig and Reactome-based gene sets and the gene set scoring mechanism is set to be released on The Comprehensive R Archive Network (CRAN).

**Fig. 1:**
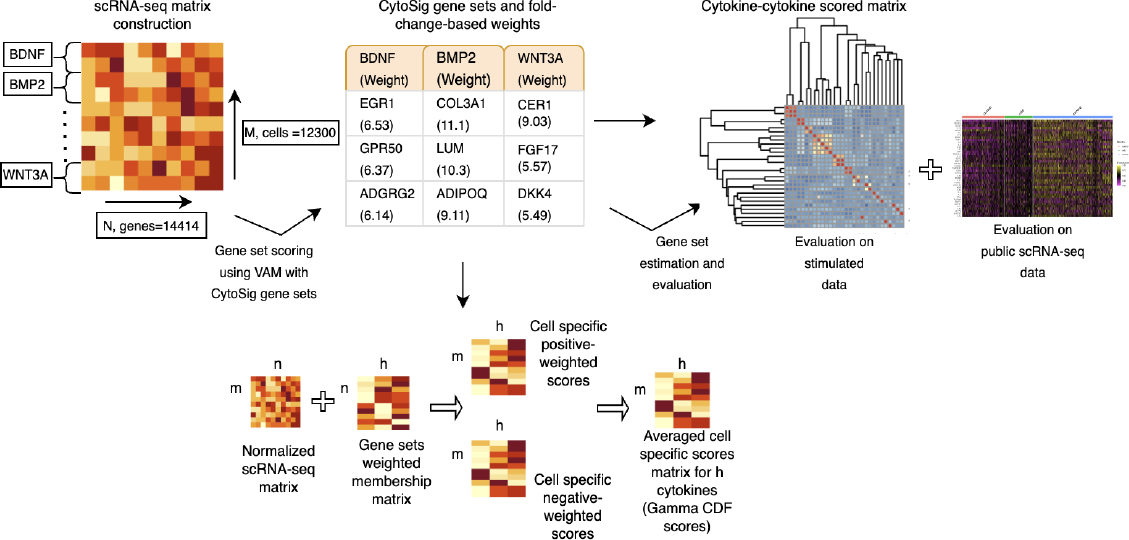
Method schematic displays the gene set construction and gene set scoring of the simulated scRNA-seq matrix containing data for up to 41 cytokines using the CytoSig database and the modified Variance-adjusted Mahalanobis (VAM) method, respectively.

### 2.2. Gene set construction

We explored two approaches for creating gene sets that capture cytokine-specific activation signatures. The first approach leveraged empirical cytokine stimulation data from the CytoSig database^11^ and the second leveraged curated cytokine signaling pathway information from the Reactome database.^12^

#### 2.2.1. CytoSig-based gene sets

CytoSig aims to simplify examination of cytokines by providing a database of target genes modulated by 43 human cytokines with bulk transcriptomics data integrated from sources such as the Sequence Read Archive (SRA),^13^ and the European Nucleotide Archive (ENA)^14^ as well as automatic extraction of the Microarray data from the Gene Expression Omnibus (GEO)^15^ and ArrayExpress (AE).^16^ For the gene set construction step, we downloaded data that captures the differential expression signature upon treatment for each cytokine from the CytoSig database. We then sorted this data by the associated log_2_ fold change (logFC) values and subset data to select for genes with positive and negative logFC values for each cytokine with gene weights set to the logFC value.

For the CytoSig-based cytokine selection, we included 41 out of the 43 cytokines included in the CytoSig database. We did not include IFN1 and interleukin-36 (IL36) in the CytoSig-specific analysis since corresponding targets for these ligands were not present in the CytoSig database version downloaded from the CytoSig application.

#### 2.2.2. Reactome-based gene sets

Reactome is a peer-reviewed, open-source and manually curated pathway database. For the gene set construction step, we utilized the Reactome database by automatically parsing the XML pathway files for the signaling pathways associated with each cytokine to retrieve molecules contained in pathways downstream of each cytokine. For example, for interleukin-2 (IL2), we extracted molecules for the ‘RAF/MAP kinase cascade,’ the ‘Interleukin receptor SHC’ and ‘Interleukin-2 signaling’ pathways and the ‘RUNX1 and FOXP3 control the development of regulatory T lymphocytes (Tregs)’ pathway since all four encapsulated IL2 signaling along with signature patterns from agents downstream of IL2.

For the Reactome-based cytokine selection, we referenced the Ramilowski et al.^17^ database to construct a ligand to receptor mapping. Amongst the 43 cytokines originally referenced in the CytoSig database, we removed IFN1 and IL36 since corresponding targets for these ligands were not present in the CytoSig database. Of the remaining 41 cytokines, we further excluded eight since we could not locate a specific matching analog for these in the Ramilowski database. These cytokines included granulocyte colony stimulating factor (GCSF), granulocyte macrophage colony stimulating factor (GMCSF), tumor necrosis factor alpha (TNFA), type-III interferons (IFNL), interleukin-12 (IL12), nitrosyl ligand (NO) and TNF-related apoptosis-inducing ligand (TRAIL). We additionally excluded Activin A, growth differentiation factor 11 (GDF11), lipoteichoic acid (LTA) and colony stimulating factor 1 (CSF1). Overall, we obtained matching receptors for 30 cytokines for the subsequent Reactome gene set construction step.

### 2.3 Gene set scoring

To estimate cytokine activity in a target scRNA-seq dataset, the SCAPE method performs cell-level scoring of either the CytoSig or Reactome-based gene sets for most of the supported cytokines. To address the need for effective gene set analysis of single cell data, our lab has previously developed both the Variance-adjusted Mahalanobis (VAM)^18^ and Reconstruction Set Test (RESET)^19^ methods. These techniques enable computationally efficient cell-level gene set scoring that can account for the sparsity and noise of single cell transcriptomic data and effectively detect patterns of differential expression and differential covariance. For the SCAPE method, we choose to use a modification of the VAM technique since it has better performance, relative to RESET, for the detection of differential expression and can take advantage of gene set weights. Specifically, for a *m × n* matrix *X*_*R*_ that holds scRNA-seq data for *m* cells and *n* genes, we first perform log-normalization to output the *m* × *n* matrix 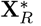 We next use VAM to compute scores for all *m* cells for each of the *h* supported cytokines using 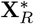 This gene set scoring generates an *m × h* matrix **S** whose elements take values between 0 and 1 and represent cumulative distribution function values for the adjusted Mahalanobis distances computed by VAM relative to a null gamma distribution.

Execution of the modified VAM method requires the following input matrices:

1. 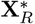 : a *m*_2_ *×n* scRNA-seq target matrix containing the positive normalized and RRR counts for *n* genes in *m*_2_ cells.
2. **A**_**p**_: a *n × h* matrix that captures the positively weighted annotation of *n* genes to *h* gene sets, i.e., gene sets for each of the *h* receptor proteins.
3. **A**_**n**_: a *n × h* matrix that captures the negatively weighted annotation of *n* genes to *h* gene sets, i.e., gene sets for each of the *h* receptor proteins.

If gene *i* is included in gene set *j*, then element *a*_*i,j*_ holds the gene weight, otherwise, *a*_*i,j*_ is not defined.

VAM outputs matrix **S**, that holds the cell-level scores for *m*_2_ cells and *h* gene sets. The computation for **S** and for *m*_2_ *× h* matrix, **M**, that holds the cell-specific squared modified Mahalanobis distances for *m*_2_ cells and *h* gene sets is detailed below (see the VAM^18^ and STREAK^20^ papers for additional detail).

#### (1) Technical variances estimation

We use the Seurat variance decomposition approach for either the log-normalized or SCTransform-normalized data^21^ to compute the length *n* vector 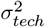 holding the technical variance of each gene in 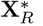 We can also otherwise set the elements of 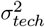 to the sample variance of each gene in 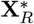 given the assumption that each gene’s observed marginal variance is entirely technical.

#### (2) Modified Mahalanobis distances computation

We compute the cell-level squared distances for a column *k* of **M**, a *m*_2_*×h* matrix of squared values of a modified Mahalanobis distance, as 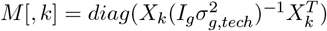. Here, *g* corresponds to the gene set (*k*) size, *X*_*k*_ is a matrix of size *m*_2_ *× g* which maps *g* columns of 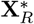 to the gene set size *k, I*_*g*_ is a *g × g* identity matrix, and 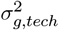 maps elements of 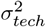 associated to the *g* genes in set *k*. We prioritize genes with large weights by dividing the elements of the 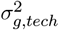 vector by the analogous elements of column *k* of matrix **A**, which aims to shrink the effective variance for genes with large weights by resulting in a larger Mahalanobis distance. We also compute modified Mahalanobis distances on a permuted version of 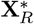 where the row labels of each column are randomly permuted. This step aims to capture the distribution of the modified Mahalanobis distances given a *H*_0_ that assumes that the normalized and RRR gene expression values in 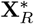 are not correlated with the technical variance.

#### (3) Gamma distribution fit

We next use the method of maximum likelihood to individually fit a gamma distribution to the non-zero elements in each column of **M**_**p**_.

#### (4) Average of cell-specific scores computation

We define the cell-level gene set scores, matrix **S**, to be the average of the gamma cumulative distribution function (CDF) value using **A**_**p**_ and one minus the CDF value usnig **A**_**n**_ for each element of **M**.

Overall, the modified VAM method sets the cell-specific scores to one minus the CDF scores, for the negative weights matrix, computed using the absolute value of the negative weights. The final cell-specific scores for matrix M are computed to be the average of the CDF scores for the positive weights matrix and one minus CDF scores for the negative weights matrix.

### 2.4. Evaluation

We evaluated the SCAPE method using both simulated and real scRNA-seq datasets that capture the transcriptional impact of stimulation by one of the 41 supported cytokines.

#### 2.4.1. Evaluation methods

- SCAPE-based CytoSig trained gene sets: Simulated scRNA-seq count matrix consisting of 12,300 total cells with 300 cells from each of the 41 cytokine conditions was estimated using the *scapeForSeurat* function from the SCAPE v0.1.0 R package with the *database* argument specified to be *cytosig* and the *normalize* parameter set to be true.
- SCAPE-based Reactome trained gene sets: Simulated scRNA-seq count matrix consisting of 12,300 total cells with 300 cells from each of the 41 cytokine conditions was subset to perform scoring using the VAM method for matched 30 cytokines (i.e., 9000 cells) using the unweighted Reactome gene sets. We performed estimation using the *scapeForSeurat* function from the SCAPE package with the *database* argument specified to be *reactome* and the *normalize* parameter set to be true.
- CytoSig: Simulated scRNA-seq count matrix consisting of 12,300 cells with data for 41 cytokines was scored using CytoSig. The generated z-scores were set as target activity scores. CytoSig prediction model of cytokine signaling activity was performed on the command line using the *CytoSig_run.py* method on Seurat counts data generated as a cell ranger output object. Normalization and log-transformation of the count data as well as mean centralization of expression values across all cells for every gene was performed as part of the CytoSig computation for the stimulated data.
- NicheNet: Cytokine activity scoring was performed using nichenetr v2.0.2 R package for 32 cytokines which overlapped with ligands in the documented ligand-receptor network. Estimation was performed using the *predict_ligand_activities* function for each one of the 32 potential ligands with each specific *geneset_oi* consisting of the marker genes differentially expressed between each of the cytokine-stimulated cell subpopulations and the remaining cells using the *FindMarkers* function from Seurat. All parameters were set to default. The area under the receiver operating characteristic (auroc) estimates was set as target activity scores. For the real COVID19 dataset, the *geneset_oi* was specified to contain differentially expressed markers between each of the specific COVID19 severity indicators (i.e., control, mild and severe) and the rest of the cells using the *FindMarkers* function from Seurat. Markers were next filtered to remove all genes with adjusted p-value greater than 0.05 and absolute log2 fold change less than 0.25, as specified in the NicheNet vignette. Lastly, the *predict_ligand_activities* function was used with all 32 ligands as *potential_ligands* and the NicheNet-specific *ligand_target_matrix* to obtain the auroc scores for the predicted ligand activities.
- Normalized RNA transcript: Simulated scRNA-seq count matrix consisting of 9,000 total cells with 300 cells from each of the 30 cytokines containing matching receptor information was scored using the analogous set of log-normalized receptor(s) gene expression data. Scoring for multiple matched receptors was performed using the VAM method. In comparison, scoring for a single matching receptor for the corresponding cytokine was performed by retaining normalized expression for that receptor. For the real COVID19 data from, in addition to the features selected via the *SelectIntegrationFeatures* function, we included all analogous receptor(s) matching each of the 30 cytokines amongst the integration features for the *anchor.features* parameter and the *FindIntegrationAnchors* function. We then estimated cytokine activity by scoring using the VAM method.

#### 2.4.2. Evaluation using simulated transcriptomics data

A key challenge for the evaluation of SCAPE is the fact that scRNA-seq cytokine stimulation datasets are not available for most of the 41 supported cytokines. To address this challenge, we adopted a strategy that used the distributional characteristics of real scRNA-seq data to simulate scRNA-seq datasets matching the expected stimulation signature for each cytokine. Specifically, we used the unstimulated scRNA-seq dataset from the Cano-Gamez et al. paper^22^ where researchers polarized Naive and Memory T cells into four T helper phenotypes. In addition to stimulating cells with anti-CD3/anti-CD28 beads to induce a state mimicking the absence of cytokines, the researchers cultured cells in the absence of stimulation from various cytokines. For our purpose, we utilized the gene correlation structure of this resting cell scRNA-seq dataset to generate correlated scRNA-seq data while preserving the individual, marginal negative binomial gene distributions. For the simulated marginal negative binomial distributions, we set the mean to the product of the estimated mean from the unstimulated data and the fold-change value associated with the target genes upon treatment by the cytokine of interest using the CytoSig database.^11^ We fixed the overdispersion parameter to 100 based on findings from the Lause et al.^23,24^ paper. Genes that were not present in the CytoSig database for the select cytokine were assigned a fold-change value of one. After generating 5,269 cells by 14,414 genes scRNA-seq datasets for each supported cytokine, we created a composite stimulated scRNA-seq dataset by selecting 300 cells from each of cytokine-specific simulated datasets, resulting in 9,000 total cells for evaluation of the 30 Reactome-based gene sets and 12,300 total cells for evaluation of the 41 CytoSig-based gene sets.

#### 2.4.3. Evaluation using public scRNA-seq data

We evaluated SCAPE using the CytoSig-based weighted gene sets on four publicly accessible scRNA-seq datasets: 1) the Cano-Gamez et al.^22^ IL4 stimulated scRNA-seq data (European Genome-Phenome Archive (EGA) EGAS00001003215), 2) the Aschenbrenner et al.^25^ IL10 blockade scRNA-seq data (GEO GSE130070), 3) the Jackson et al.^26^ IL13 stimulated data (GEO GSE145013) and 4) the Liao et al.^10^ COVID19 data (GEO GSE145926). Each dataset was individually processed using Seurat v.4.1.0^21,27–29^ in R v.4.1.2.^30^ For each of the publicly stimulated dataset, we merged across replicates and integrated across stimulation, blockade (i.e., IL4, IL10 and IL13 data) or disease severity categories (COVID19 dataset).

## 3. Results and Discussion

### 3.1. Validation on simulated scRNA-seq data

The cytokine activity predictions generated by SCAPE on the simulated scRNA-seq data can be held in a matrix where the rows represent the simulation model (i.e., the cytokine whose expected stimulation signature is simulated), the columns represent the cytokine whose activity is estimated by SCAPE, and the matrix elements contain the average estimated activity of the target cytokine across all cells in all datasets simulated according to a particular cytokine model. For SCAPE estimates generated using the CytoSig-based gene sets, this is a 41 *×* 41 matrix, as visualized in Figure 2. For SCAPE estimates generated using the Reactome-based gene sets, this is a 30 *×* 30 matrix, as visualized in Figure 3 (see Figures 4-6 for scored simulation matrix computed on the remaining comparative methods). When the ordering of cytokines for rows and columns is identical, these matrices should have a diagonal structure if the activity estimation is accurate, i.e., the cytokine used to simulate the data will be the cytokine with the largest estimated activity. For the CytoSig-based gene sets, all cytokines have average cell-level scores that align with the simulated cytokine stimulation model, a pattern that is also indicated by the distinctively higher average cell-level score expression distribution along the diagonals. In comparison, for the Reactome-based gene sets, the scored cytokine activity shows very little correspondence with the simulated model.

**Fig. 2:**
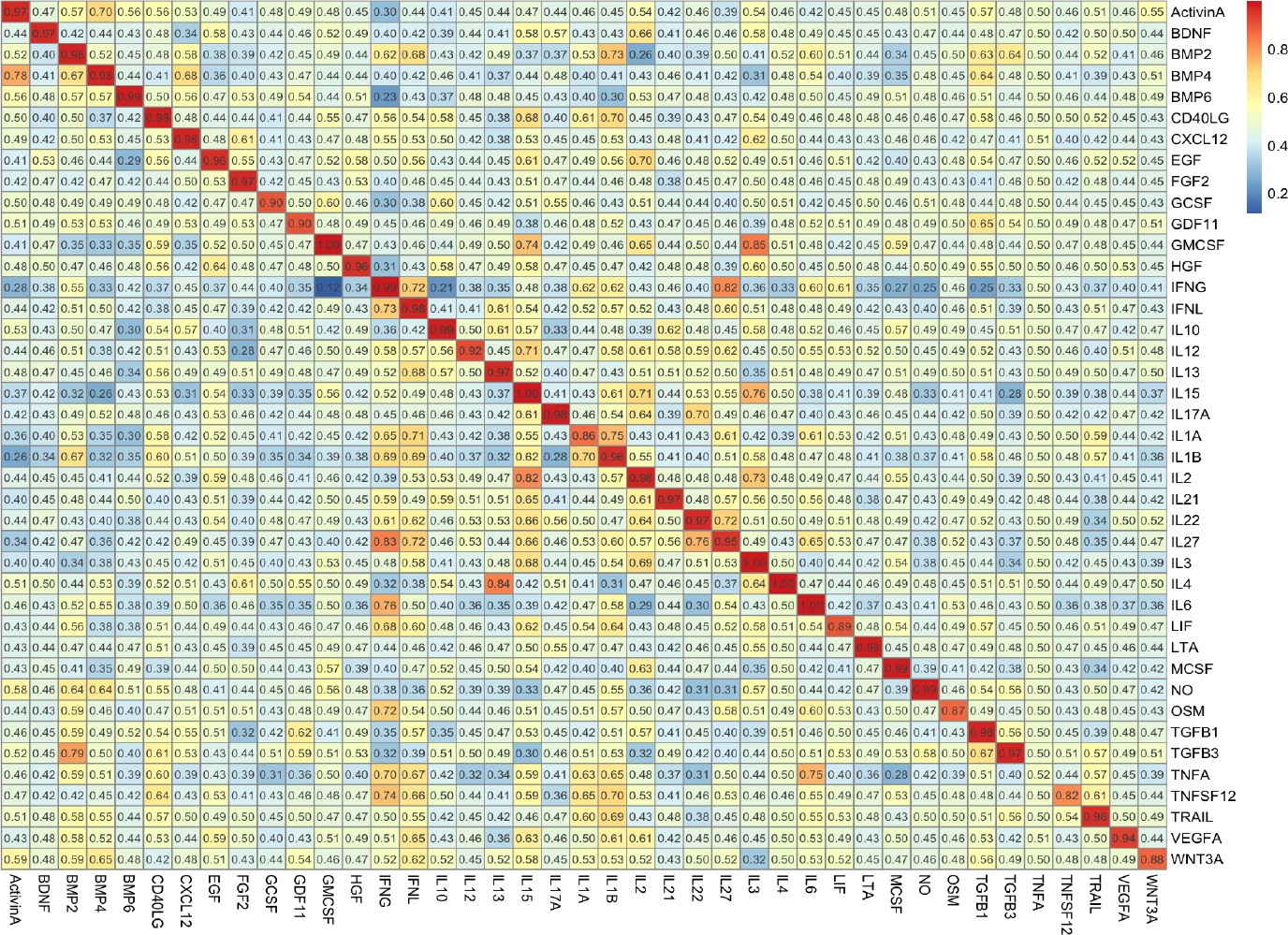
Scored simulation matrix for the SCAPE-based CytoSig trained gene sets algorithm where rows reflect the simulated cytokine stimulation, the columns reflect the target cytokine for activity estimation and the matrix elements contain the average estimated activity of the target cytokine across all cells in all datasets simulated according to a particular cytokine model.

**Fig. 3:**
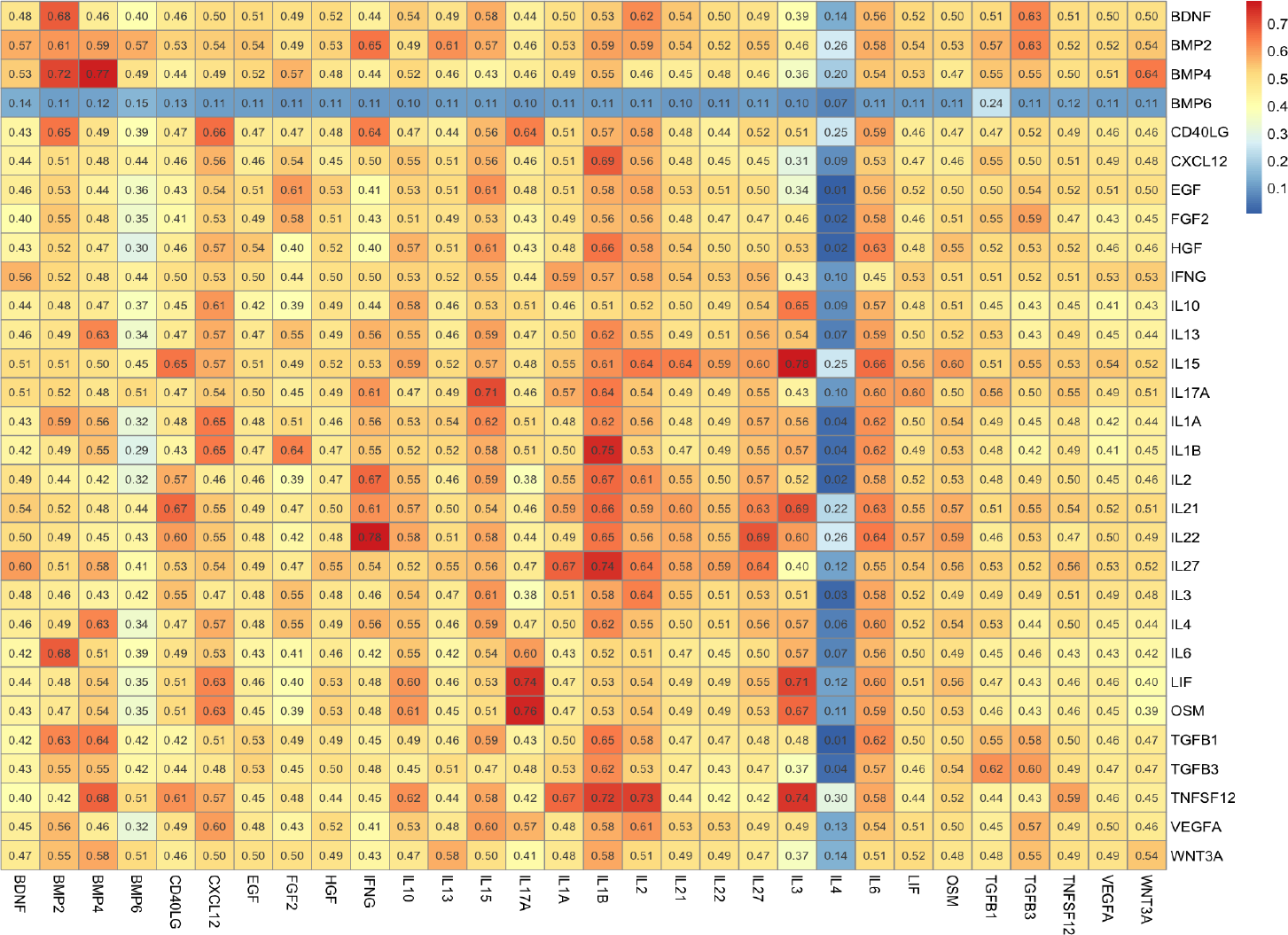
Scored simulation matrix for CytoSig-based Reactome trained gene sets.

**Fig. 4:**
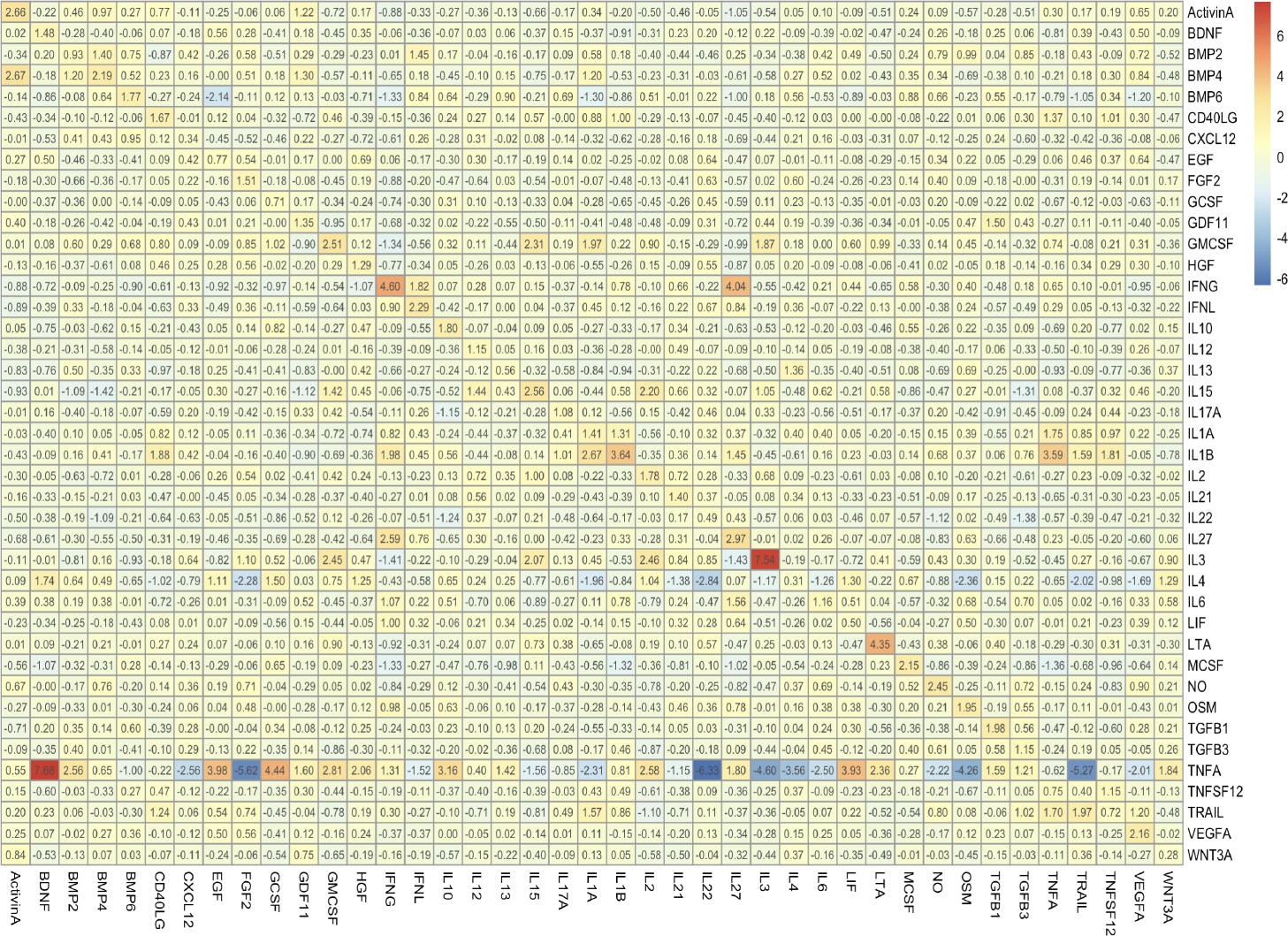
Scored simulation matrix using the CytoSig algorithm.

In addition to visualizing the mean CDF scores for each method, we computed a measure to quantify the percentage of cytokines that have correspondence between the rows and columns along the diagonals for cytokine estimates generated using the SCAPE-based CytoSig trained gene sets and comparative techniques. As observed in Figure 18, all 41 cytokines had the highest correspondence along the diagonals when estimated using SCAPE-based CytoSig trained gene sets. This compares with 30 out of 41 cytokines that have correspondence along the diagonals when estimated using the CytoSig algorithm, 13 out of 32 for cytokines estimates using NicheNet and 3 cytokines out of 30 when estimated using the Reactome and the RNA normalized algorithms.

### 3.2. Validation on real public stimulated and blockade experiment data

Following validation on the simulated dataset, we next validated the SCAPE method using the CytoSig-based gene sets on individual stimulation and blockade experiment datasets indicated in Table 1. For each data, we separately normalized and integrated unstimulated and stimulated data for the IL4 cytokine, acute stimulated or chronic stimulated IL13 data or stimulated and blockade IL10 data. We then implemented the *scapeForSeurat* function for each dataset. We assessed the effectiveness of our method by computing the mean CDF scores on a non-log scale by exponentiating before the mean computation. We hypothesized that a higher average CDF score following regular or chronic stimulation and a lower average CDF score following blockade experiment will capture up or downregulation activity signal from the stimulation and blockade experiments, respectively. As detailed in Table 2, differences between the mean estimated activity of the target cytokine in the perturbed versus control conditions were statistically significant for all three analyzed datasets at a type I error rate of 0.05. Table 2 also reports the corresponding standard deviation and 95% confidence intervals, as computed using the Wilcoxon rank-sum test.

**Table 1:**
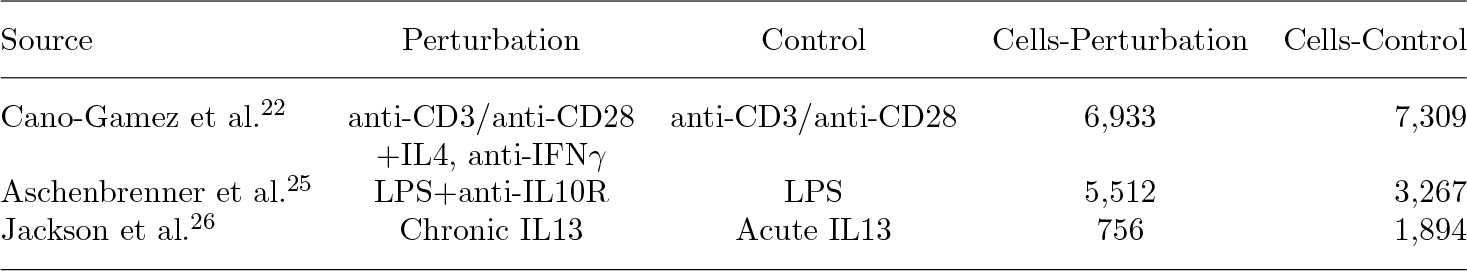
Public single-cell transcriptomics datasets used for validation. The ‘Cells-Perturbation’ and ‘Cells-Control’ columns indicate the number of cells used in each condition.

**Table 2:** Assessment of median and interquartile range (IQR) for differences in predicted activity scores between the perturbed and the control states for each cytokine condition with confidence intervals computed on differences in the predicted activity scores between the perturbed and the control states using the Wilcoxon rank-sum test.

| Perturbation | Perturbation Mean (SD) | Control Mean (SD) | 95% Confidence Interval [p.value] |
| --- | --- | --- | --- |
| IL4<sup>22</sup> | 0.757 (0.746, 0.768) | 0.630 (0.617, 0.642) | (0.104, 0.138) [$< 0.0001$] |
| IL10<sup>25</sup> | 0.606 (0.597, 0.616) | 0.894 (0.880, 0.908) | (-0.300, -0.265) [$< 0.0001$] |
| IL13<sup>26</sup> | 0.762 (0.731, 0.792) | 0.697 (0.676, 0.717) | (0.0359, 0.111) [$< 0.0001$] |

We first validated the SCAPE IL4 activation estimates using the Cano-Gamez et al. dataset, where researchers stimulated activated T cells with anti-CD3/anti-CD28 to generate an effector phenotype (Th0 state) that was subsequently stimulated by different cytokines, including IL4 (i.e., the Th2 state stimulated with anti-CD3/anti-CD28 and IL4). We report mean IL4 activity score of 0.757 (0.746, 0.768) for the Th2-IL4 stimulated/perturbed state and a mean of 0.630 (0.617, 0.642) for the Th0-T cell activated unstimulated/control state (95% CI for the mean difference was 0.104 to 0.138). The reported mean activity score for IL4 stimulated state is not only higher than the unstimulated state, it is also higher than the average predicted cytokine activity for all other stimulated cytokines, as indicated by the Stim-IL4 column in Figure 7.

**Fig. 5:**
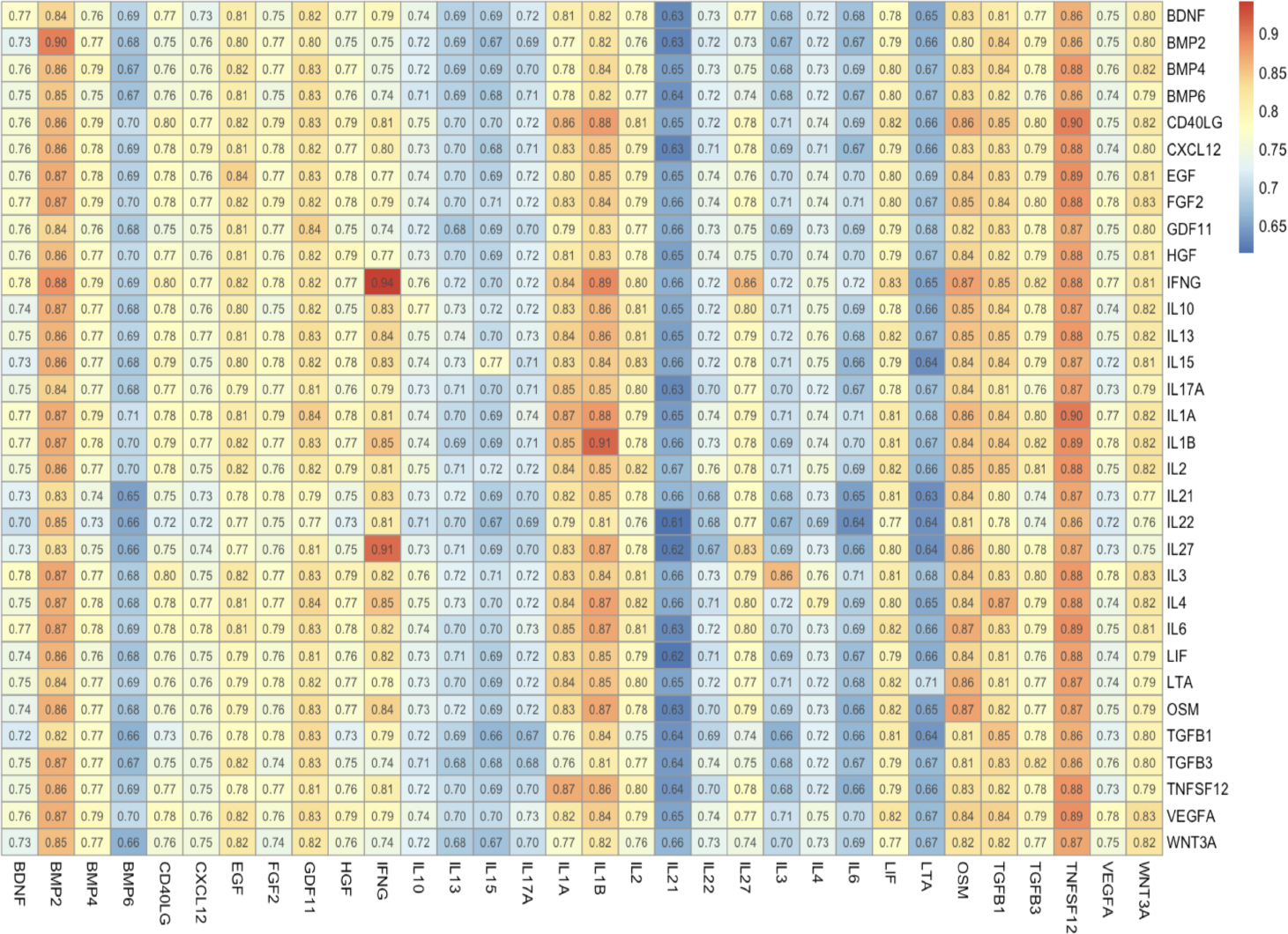
Scored simulation matrix using the NicheNet ligand prediction framework.

**Fig. 6:**
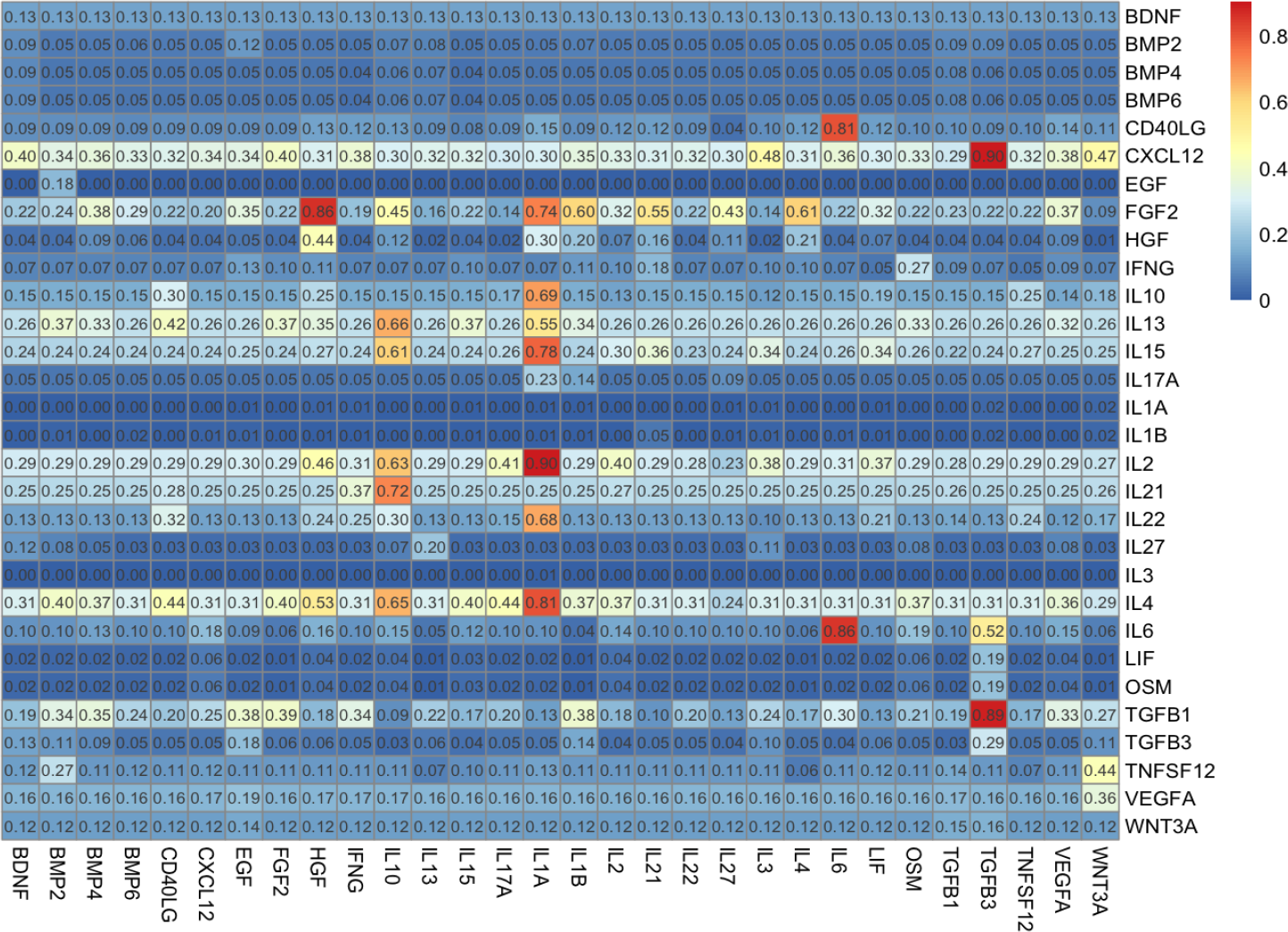
Scored simulation matrix using the RNA normalized algorithm.

**Fig. 7:**
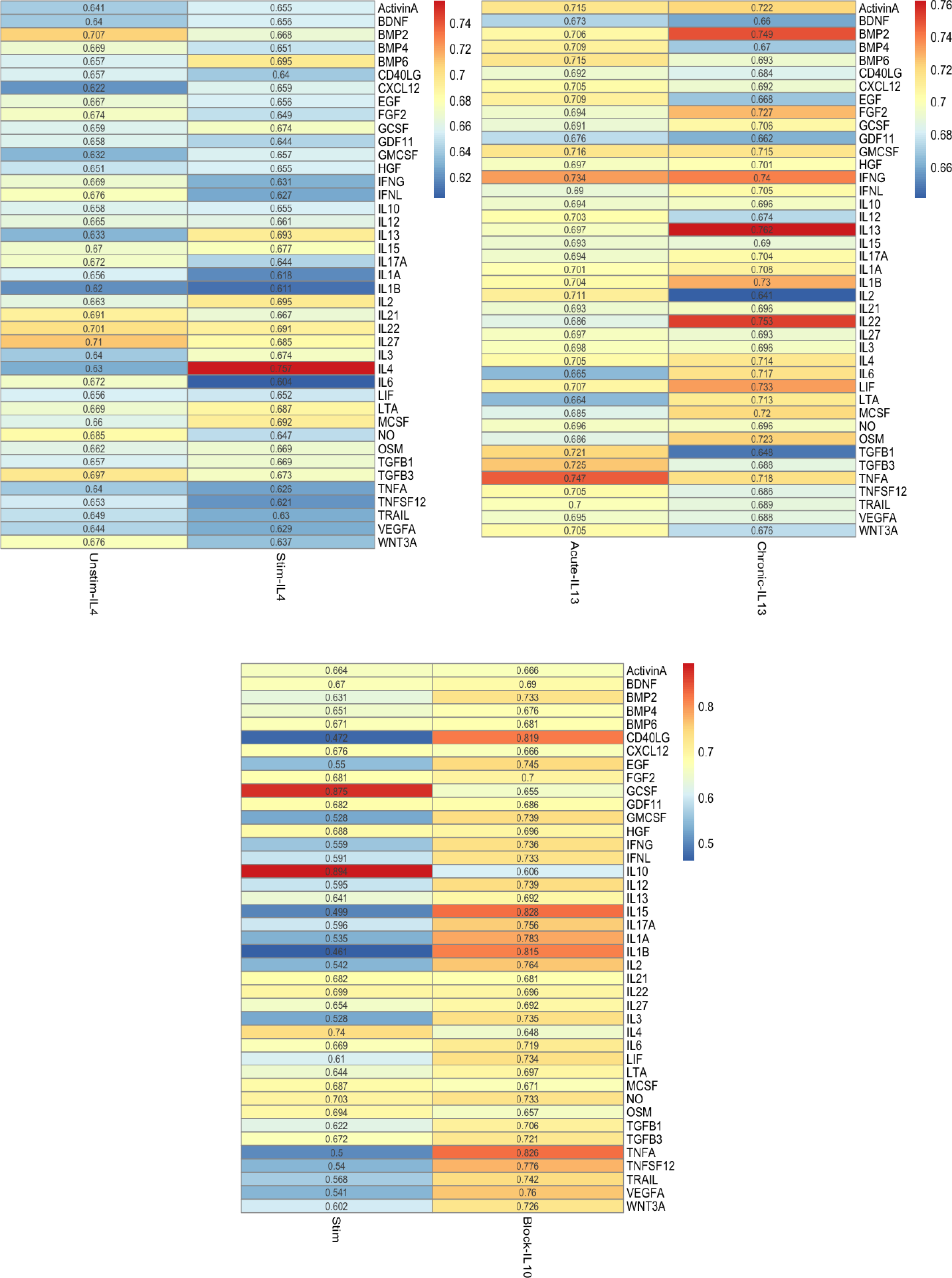
Average predicted cytokine activity scores for the IL4 stimulation experiment condition (top left), IL13 stimulation condition (top right) and IL10 blockade condition (bottom).

We next validated the SCAPE IL13 activation estimates using the Jackson et al.^26^ dataset, where researchers stimulated human airway epithelial cell cultures with acute and chronic IL13. We report mean activity score of 0.762 (0.731, 0.792) for chronic IL13 stimulation and mean activity score of 0.697 (0.676, 0.717) for acute IL13 stimulation (95% CI for the mean difference was 0.0359 to 0.111). Similarly as with IL4, the reported mean activity score for chronic IL13 stimulated state is not only higher than the acute IL13 stimulated state, it is also higher than the average predicted cytokine activity for all other chronic stimulated cytokines, as indicated by the Chronic-IL13 column in Figure 7.

Lastly, we validated the SCAPE IL10 blockade estimates using the Aschenbrenner et al. study.^25^ In that work, the authors examined markers for inflammatory bowel disease using LPS stimulation with or without concomitant IL10 signaling blockade. We report mean activity score of 0.606 (0.597, 0.616) for the IL10 blockade category and mean activity score of 0.894 (0.880, 0.908) for the LPS stimulation category (95% CI for the mean difference was −0.300 to −0.265). We report the mean activity scores for IL10 blockade to be lower than the average activity scores for all other cytokines, as displayed in the Block-IL10 column in Figure 7. Additionally, we note that the decrease in IL10 activity coincides with an increase in the mean predicted activity scores for GMCSF (0.739 for LPS+anti-IL10R vs. 0.528 for LPS), IL1A (0.783 for LPS+anti-IL10R vs. 0.535 for LPS), IL12 (0.739 for LPS+anti-IL10R vs. 0.595 for LPS), and IFN*γ* (0.736 for LPS+anti-IL10R vs. 0.559 for LPS) cytokines. This result is supported by the Aschenbrenner et al. study as researchers note that the addition of IL10 blockade to LPS significantly upregulates LPS-induced factors such as IL23p19, GM-CSF, IL-1*α*, IL-12p70, IL-12p40 and IFN*γ*. Overall, these comprehensive activity profiles not only validate the magnitude and direction of the target cytokine for each experiment, but also indicate that the SCAPE method based on CytoSig trained gene sets can accurately recapitulate biological heterogeneity related to real, transcriptomics-based stimulation and blockade models.

### 3.3. Validation on real public COVID19 data

In addition to validation on real stimulated and blockade experimental data, we estimated SCAPE and comparative techniques on the Liao et al. COVID19 dataset, which contained data for three control, three mild and six severe patients. We merged data across replicates and integrated across disease severity categories using Seurat’s reciprocal PCA (‘RPCA’) algorithm.^21,27–29^ Since normalization was performed on each dataset before integration, the integrated data was not normalized when performing estimation using the *scapeForSeurat* function. The estimation was similarly performed on the integrated assay for all comparative techniques.

We assessed the predicted cytokine activity for the COVID19 dataset by computing the mean CDF scores on a non-log scale. In this case, we hypothesized to observe a relative increase in the average scores from the control to mild to severe COVID19 categories. While this approach may not apply to all cytokines, we think it will apply to the majority and allow us to capture the graded response to the cytokine signaling influx in response to COVID19. There is further evidence for this hypothesis from Del Valle et al.^31^ who reported that levels of cytokines such as IL-6 and interleukin-8 (IL-8) correlated with COVID19 severity (i.e., progressing from moderate, severe and severe with end organ damage) taking into account metrics such as lung imaging and creatinine clearance. In addition to the heatmaps that displayed the raw estimated scores, as displayed in Figures 8-11 for the SCAPE-based CytoSig trained gene sets, and the average predicted cytokine activity scores, as displayed in Figures 16 and 17, we computed a metric to quantify the number of cytokines that both displayed a statistically significant overall trend as well as showed an increasing average trend given disease progression from control to mild to severe COVID19. We performed the non-parametric Kruskal-Wallis test to inspect for overall trend differences. We display the overall trends in the predicted cytokine activity for each cytokine over the COVID19 severity category for all methods in Figures 12-15. Bold text label is used to indicate cytokines which fulfill both criteria (i.e., a statistically significant trend and a positive increase from control to mild and from mild to disease categories). Lastly, we summarize the proportion of cytokines that have a statistically significant trend, at an alpha level of 0.05, and that show an increasing positive trend in the average CDF scores from control to mild to severe COVID19 disease type for all comparative methods, including SCAPE-based CytoSig trained gene sets, the CytoSig algorithm, SCAPE-based Reactome trained gene sets and the RNA-normalized algorithm in Figure 18. In this case, NicheNet was not chosen as a comparative method since it does not generate single-cell-level estimates of predicted ligand activity but only generates the ranking of the ligands.

**Fig. 8:**
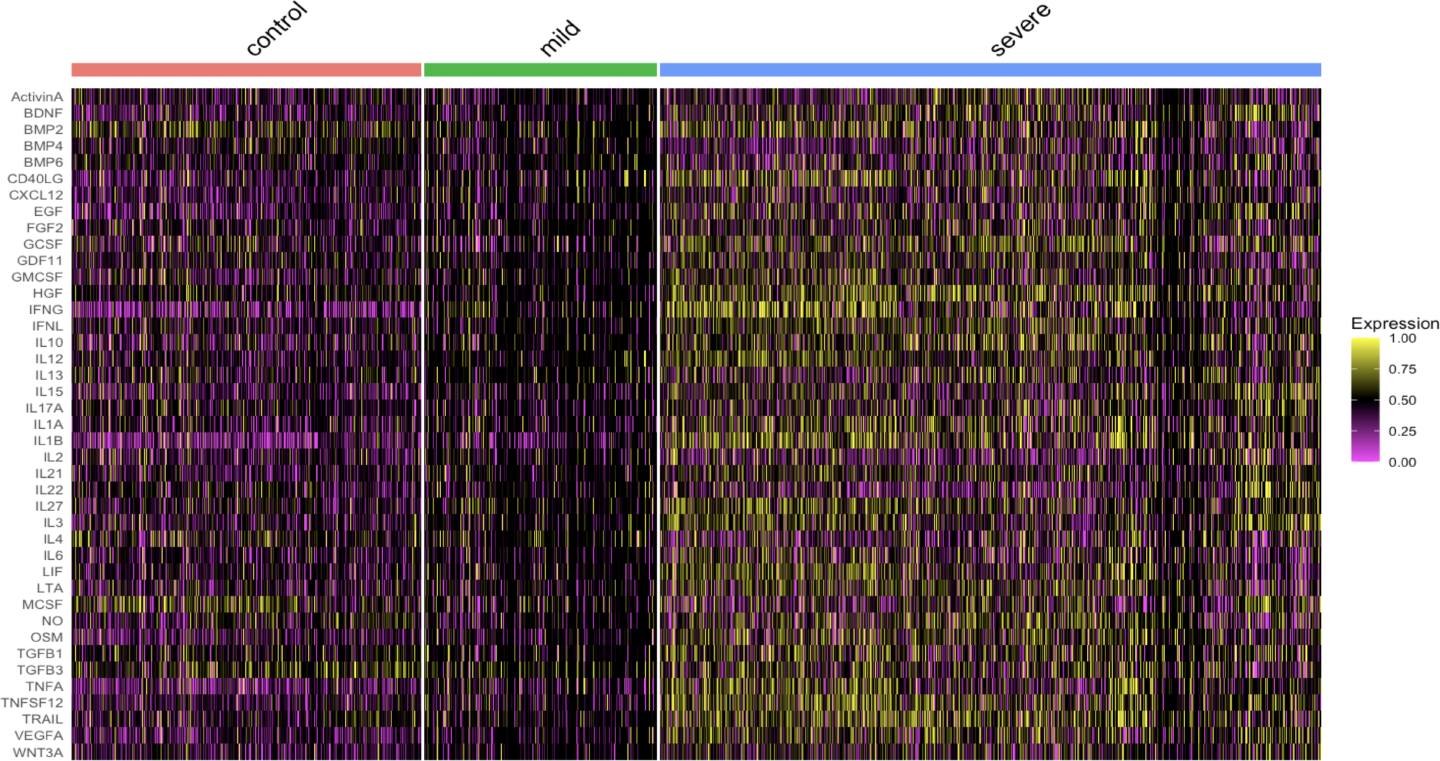
Application of SCAPE-based CytoSig trained gene sets to the COVID19 dataset and a visualization of the estimated cytokine activity scores.

**Fig. 9:**
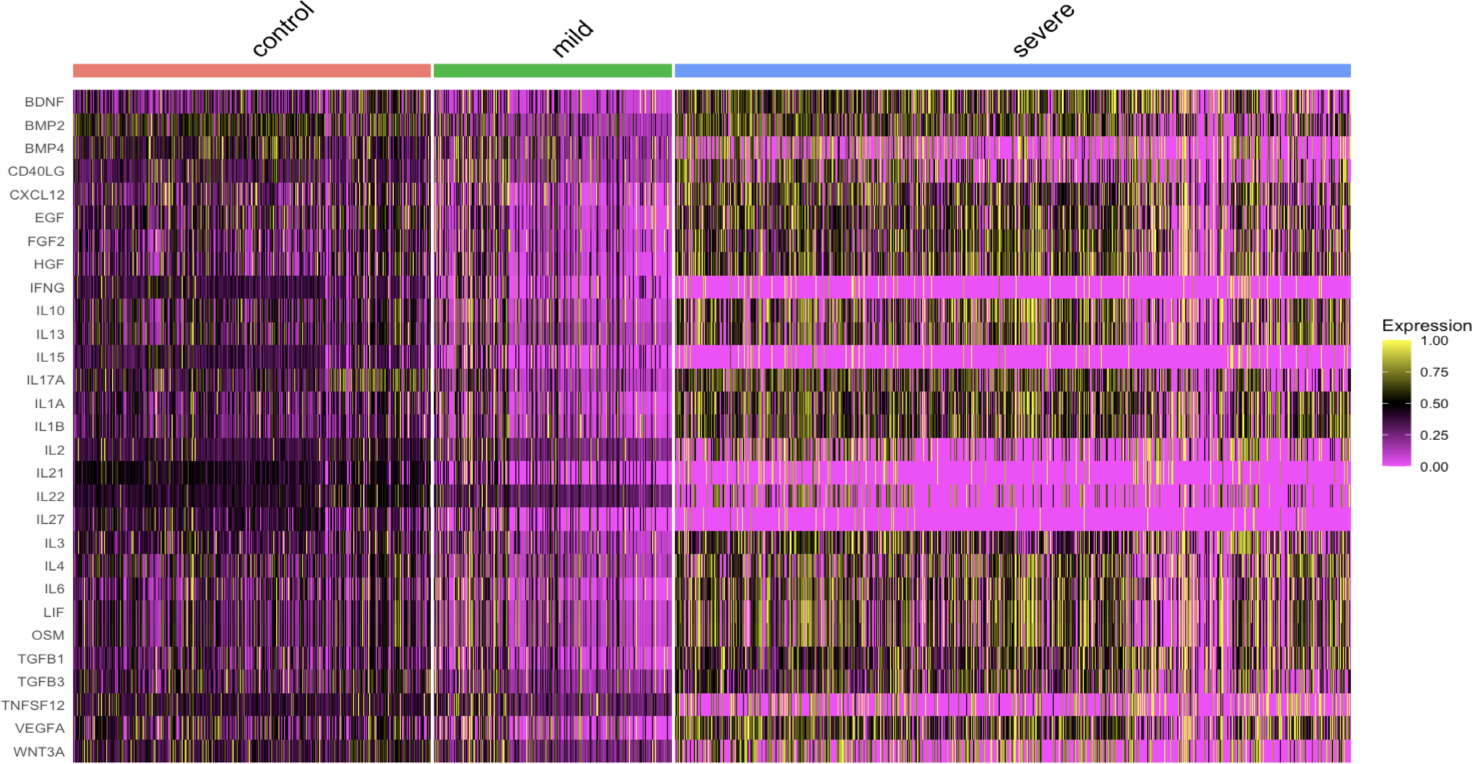
Application of SCAPE-based Reactome trained gene sets to the COVID19 dataset.

**Fig. 10:**
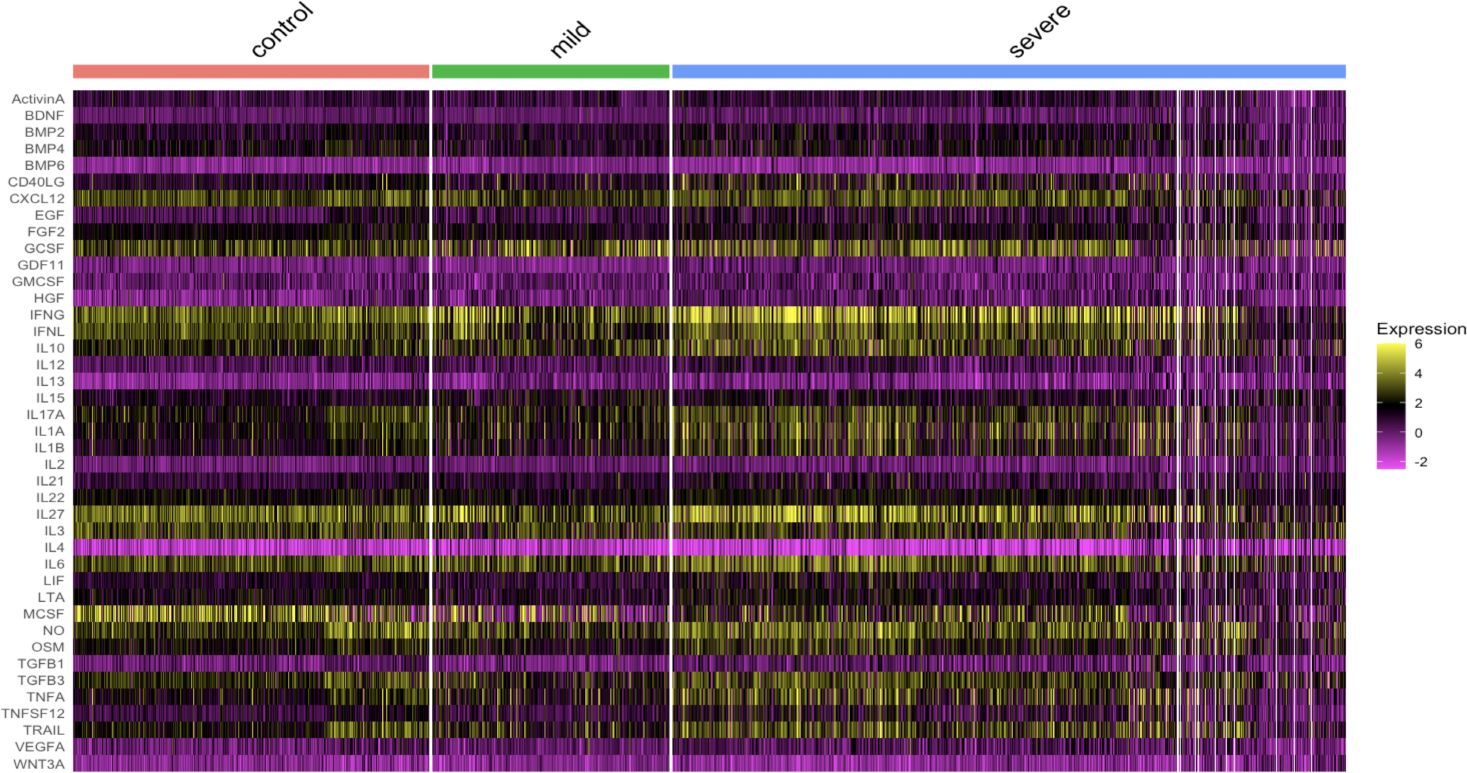
Application of CytoSig to the COVID19 dataset.

**Fig. 11:**
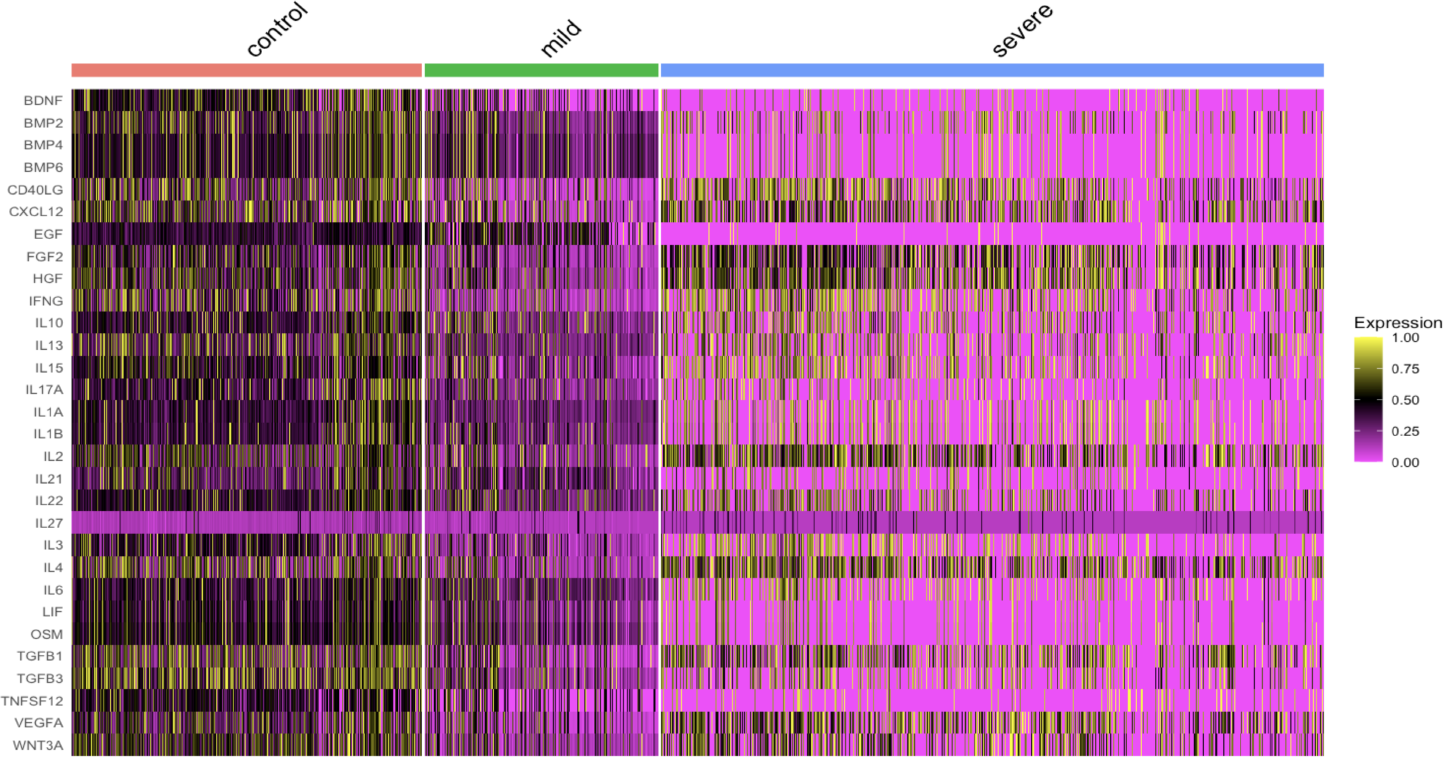
Application of RNA Normalization algorithm to the COVID19 dataset.

**Fig. 12:**
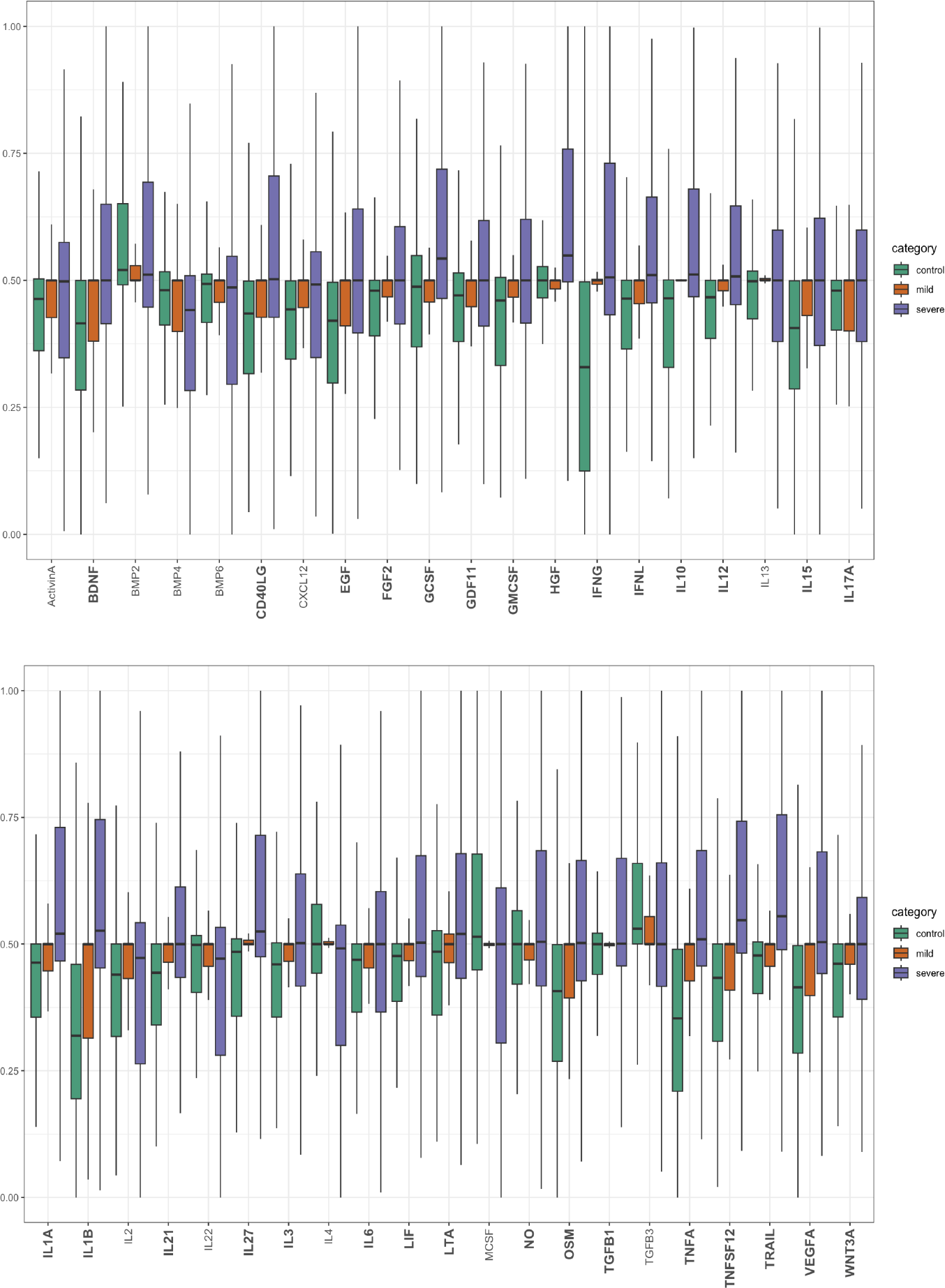
COVID19-based boxplots for SCAPE-based CytoSig trained gene sets. Bold cytokines have a statistically significant difference at an alpha level of 0.05 (based on the non-parametric Kruskal Wallis test) and indicate presence of a continuously increasing positive trend from the control to mild to severe COVID19 diseased patients.

**Fig. 13:**
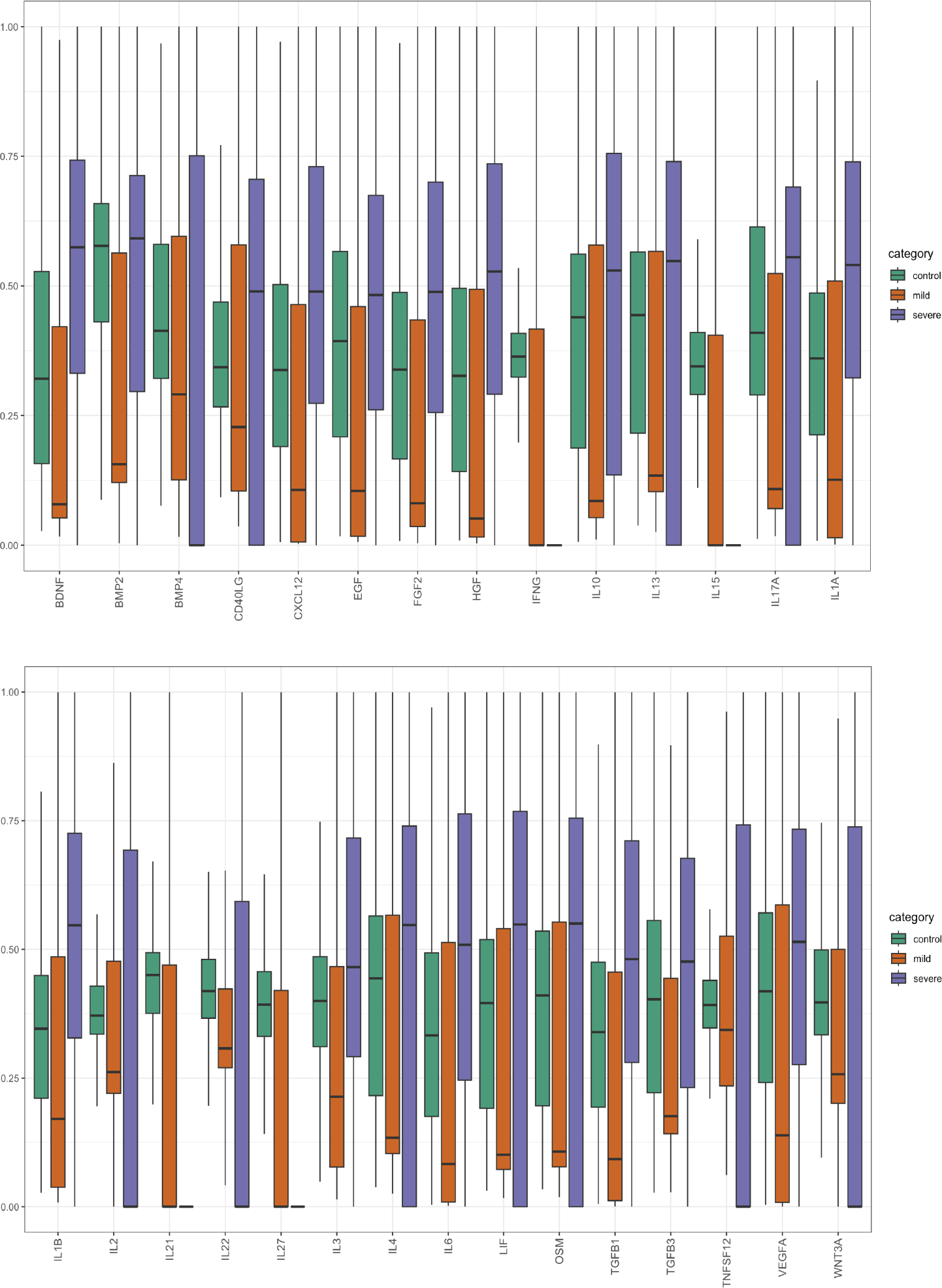
COVID19-based boxplots for SCAPE-based Reactome trained gene sets.

**Fig. 14:**
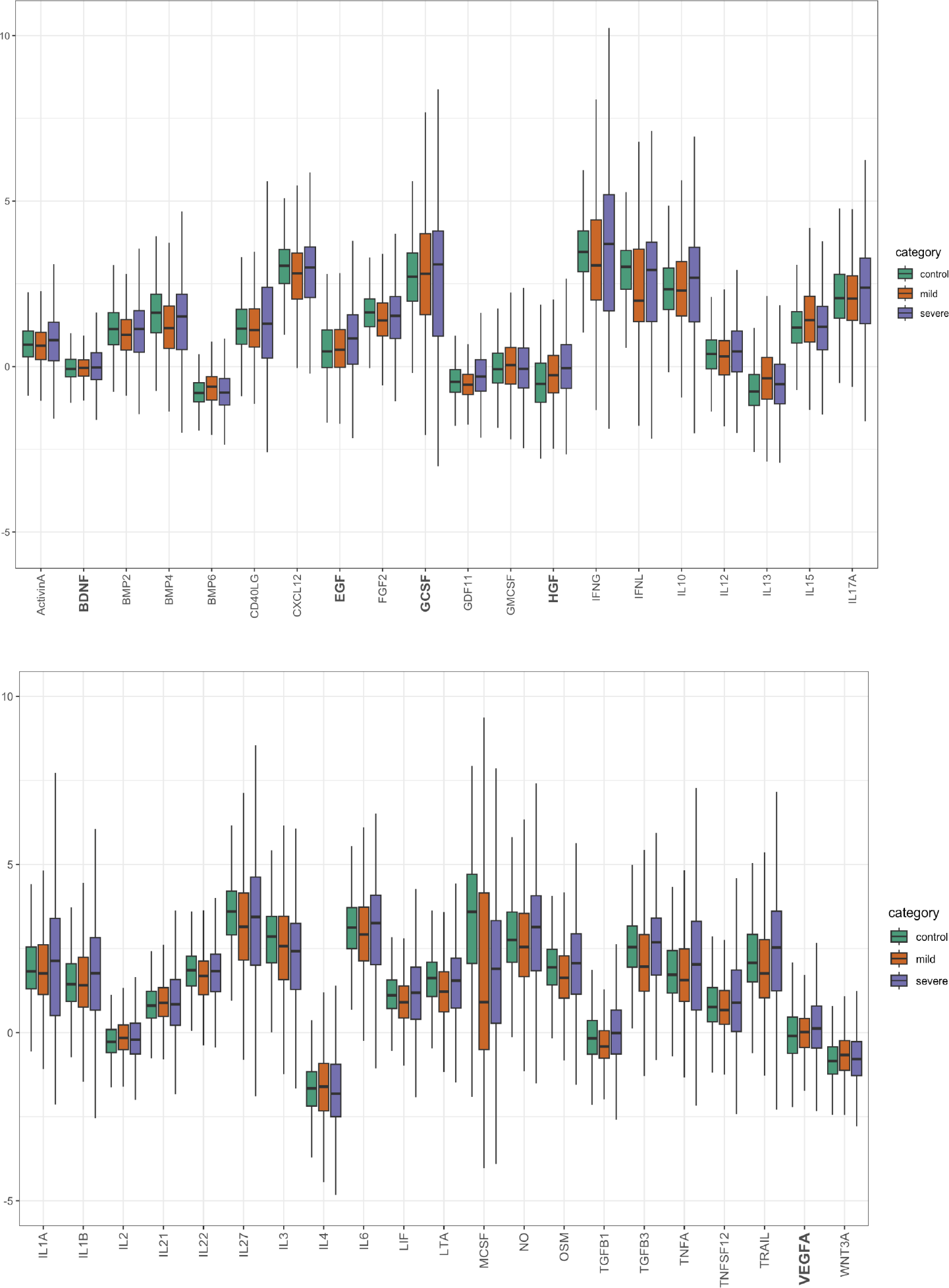
COVID19-based boxplots for the CytoSig algorithm.

**Fig. 15:**
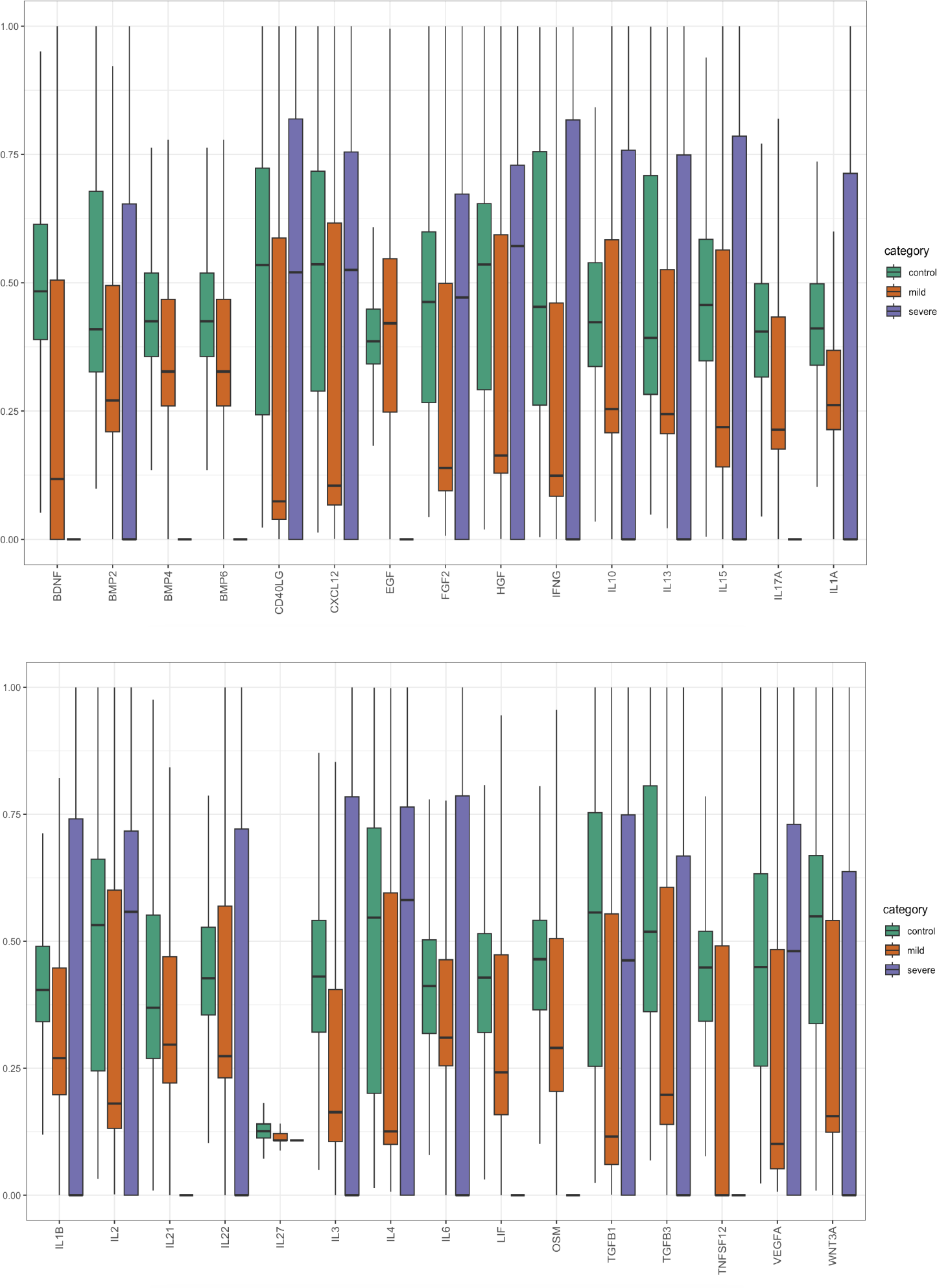
COVID19-based boxplots for the RNA-normalized algorithm.

**Fig. 16:**
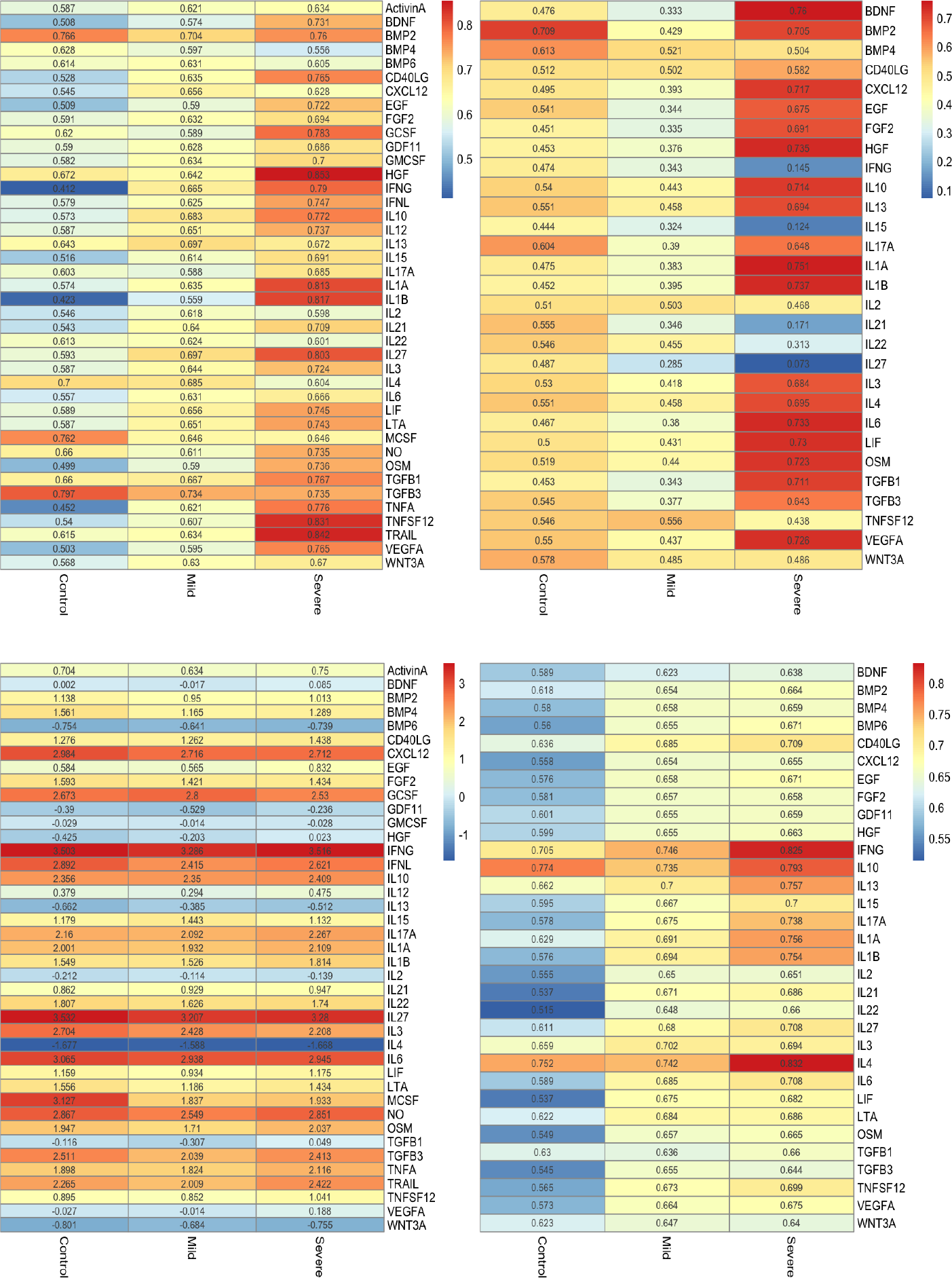
Average of predicted cytokine activity scores for the SCAPE-based CytoSig trained gene sets (top left), SCAPE-based Reactome trained gene sets (top right), the CytoSig algorithm (bottom left) and the NicheNet ligand prediction framework (bottom right).

**Fig. 17:**
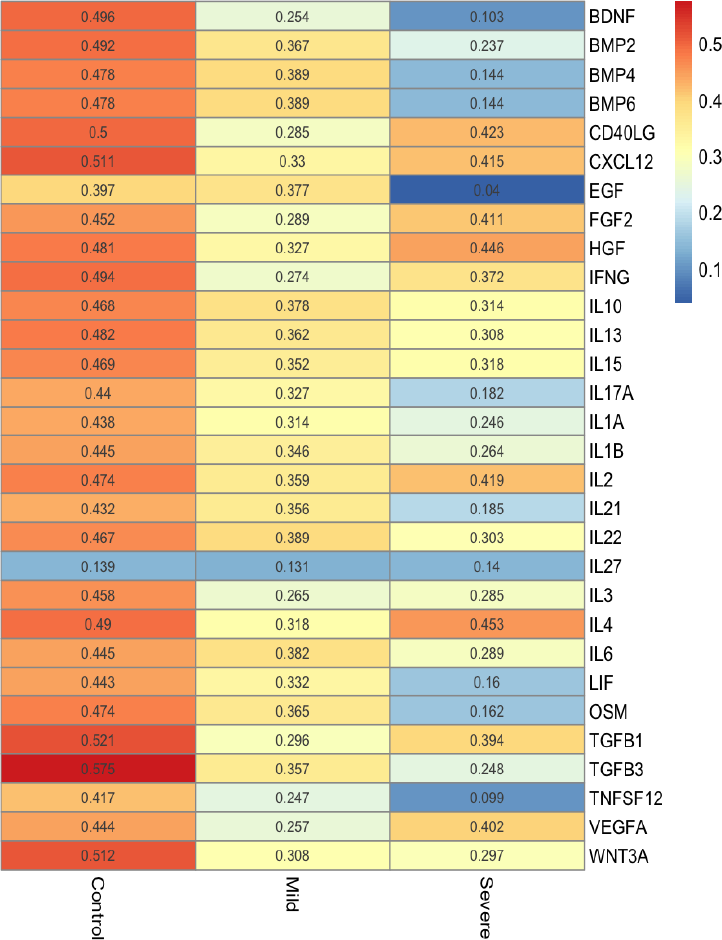
Average of predicted cytokine activity scores for the RNA-normalized algorithm.

**Fig. 18:**
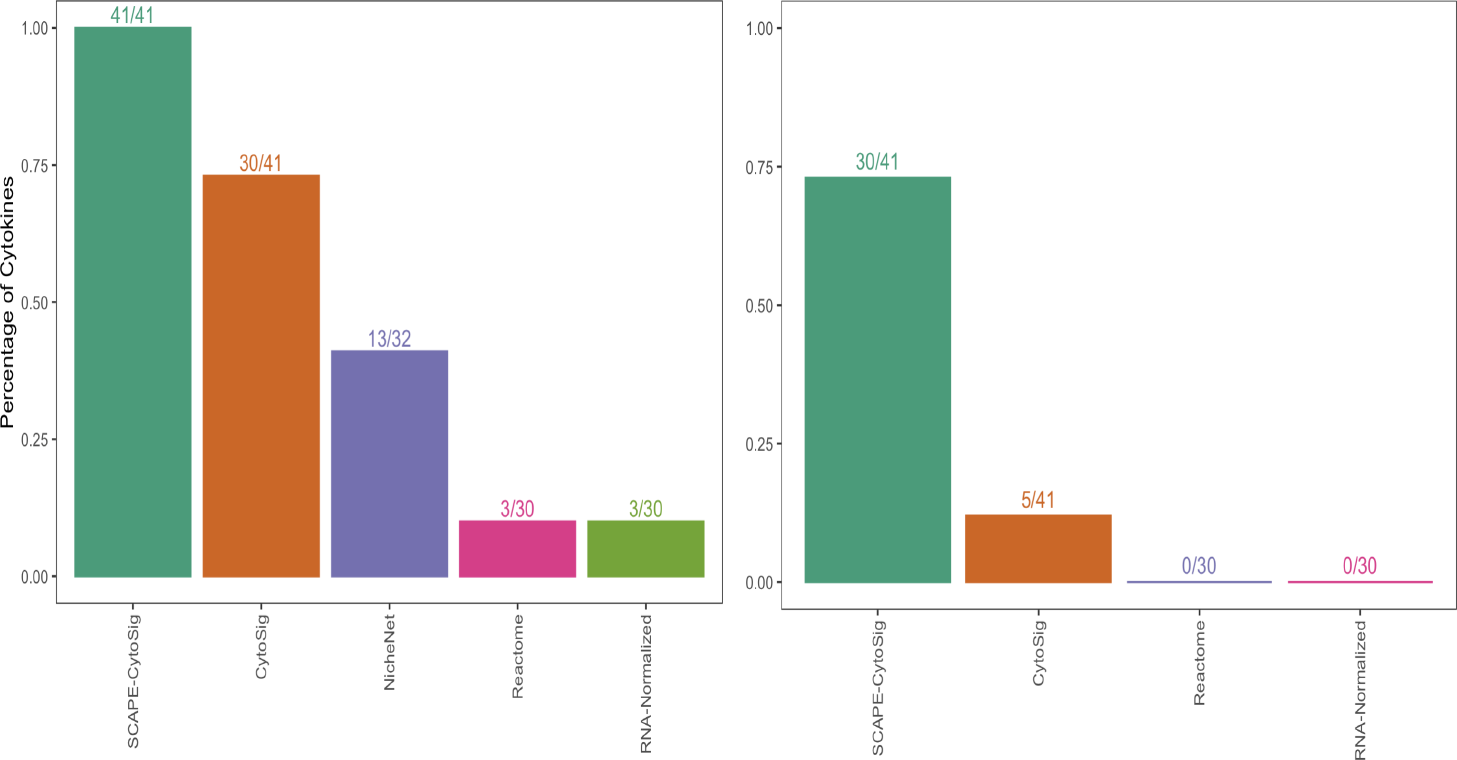
Proportion of cytokines that have the highest score along diagonal for the simulated data for each comparative method (left) and proportion of cytokines with a progressively positive trend of increasing average predicted cytokine activity scores from control to mild to severe COVID19 type that are additionally statistically significant at an alpha level of 0.05 (based on the Kruskal Wallis test) (right).

Overall, as displayed in Figure 18, we observe that the estimates generated by SCAPE-based CytoSig trained gene sets have the highest proportion of cytokines (30/41) that show the progressively increasing trend of average CDF scores over COVID19 severity as compared to the estimates generated by the CytoSig, Reactome and the RNA-normalized algorithms. We observed this positive trend progression for IL6 and for additional inflammatory cytokines such as IL1*β* and TNF that are known to be associated with COVID19 disease severity.^10^

## 4. Conclusion

In this paper, we introduce the SCAPE (Single cell transcriptomics-level Cytokine Activity Prediction and Estimation) method for cell-level cytokine activity analysis of scRNA-seq data.

SCAPE performs activity estimation by scoring weighted cytokine-specific gene sets that are created using gene-level associations learned from the CytoSig database. As demonstrated by evaluations on both simulated and real scRNA-seq data, the SCAPE method generates estimates that match expected cytokine activity. The SCAPE method has practical utility for precision medicine applications where researchers or clinical practioners need to accurately quantify cytokine activity at the level of individual cells within a tissue sample. Such applications have particular relevance for the study of cytokine-related risk factors associated with conditions such as autoimmune disease, viral infection, and cancer immunotherapy.

